# vHULK, a new tool for bacteriophage host prediction based on annotated genomic features and deep neural networks

**DOI:** 10.1101/2020.12.06.413476

**Authors:** Deyvid Amgarten, Bruno Koshin Vázquez Iha, Carlos Morais Piroupo, Aline Maria da Silva, João Carlos Setubal

## Abstract

The experimental determination of a bacteriophage host is a laborious procedure. For this reason, there is a pressing need for reliable computational predictions of bacteriophage hosts in phage research in general and in phage therapy in particular. Here, we present a new program called vHULK for phage host prediction based on 9,504 phage genome features. These features take into account alignment significance scores between predicted-protein sequences in the phage genomes and a curated database of viral protein families. The features were fed to a deep neural network, and four distinct models were trained to predict 61 different host genera and 52 host species. In random controlled test sets, the program obtained 99% and 98% accuracy values at the genus and species levels, respectively. On a validation dataset with 2,178 phage genomes, mean accuracies were 82% and 52% at the genus and species levels, respectively. When compared against other phage host prediction programs on the same validation dataset, vHULK achieved substantially better performance, therefore demonstrating that the program is an advance on the state-of-art in phage host prediction. vHULK is freely available at https://github.com/LaboratorioBioinformatica/vHULK.

## INTRODUCTION

Viruses are believed to be by far the most diverse and abundant biological entities in the biosphere (1). They are present in virtually any place or environment as long as a host is nearby. Acquisition of new knowledge about viruses has been restricted by our capacity of sampling the environment and isolating new strains through cultivation-based methods. Recent improvements in Next Generation Sequencing techniques and the rise of metagenomics as a prominent field of study have given us access to unprecedented genetic diversity. Researchers now have at their disposal thousands of DNA sequence datasets from different types of environmental or host-associated samples, such as ocean, soil, and animal guts (2–4). These datasets are helping us to shed light on the so-called viral dark matter (5–7). Recently, there have been efforts to consolidate many of these datasets in general databases (8,9), thereby facilitating investigation of the viral taxonomic and genomic diversity, as well as virus’ roles in specific habitats (10).

Bacteriophages (or simply phages) are a substantial part of the global virome, and as such their genomes are also being sampled at a rapid pace (6,11). Phages have achieved prominence in the last few years thanks to phage therapy, which is seen by many as a viable resource against multidrug resistant bacteria (12). A key step in phage therapy development is finding a phage that can infect and efficiently terminate with high specificity an infectious agent of interest. Establishing a link between phage and host is however far from trivial, since phages may present a variety of infecting molecular mechanisms, and a host range that varies from narrow (strain-specific) to very broad (hosts from a same genus, family or higher taxonomic levels) (13–15). Furthermore, host range characterization mostly depends on laborious wet-lab experimental methods and may be very specific for species or even bacterial strains. As examples of such techniques, we may cite the use of fosmids, viral tagging with fluorescent die, and phageFISH (16–18). Because of the laboriousness and specificity of these methods, effective computational prediction of phage hosts should be of great help.

R. Edwards and colleagues have reviewed genomic signals used to link a phage to its host (19). Oligonucleotide frequency profiles, sequence similarity, and CRISPR-spacers matches were evaluated. Sequence similarity between phage and host genome, as well as matches of CRISPR-spacers were found to be more effective in establishing phage-host pairs. However, only 40% accuracy was achieved with these features. Since then, a several other tools have become available. Tools such as Phaster (20), PhiSpy (21), Phage_Finder (22), and Prophinder (23) focus on finding phages integrated in a host genome (prophages). Despite establishing a clear phage-host relationship, these approaches are restricted to temperate phages that are integrated in their host genome at the moment of sampling.

More general approaches have been suggested, such as the use of Markov models based on *k*-mer frequencies implemented in the WiSH tool (24). According to its authors, WiSH presents 63% of mean accuracy when predicting host genus among 20 possible genera. A more recent work proposed the use of neural networks based on an ensemble of distance measurements and other features, concepts that were implemented in the tool VirHostMatcher-net (VHM-net) (25). VHM-net achieved accuracies that varied from 43% to 59% at the host genus level only. VHM-net is theoretically able to predict thousands of different prokaryote hosts. The most recent program in this field is Random Forest Assignment of Hosts (RaFAH), which has been shown to outperform previous virus-host prediction methods at the phylum level (26).

In this work, we present a novel approach for phage host prediction based on a unique representation of annotated genomic features and deep neural networks. Features are extracted from phage genomes and searched against a protein database, which provides information about genes and their function. These features are converted to a pixel-like matrix and fed to multi-layer perceptron models (MLP) trained to predict 52 and 61 different host species and host genera, respectively. Trained models were implemented in a user-friendly tool named vHULK, which stands for Viral Host UnveiLing Kit. vHULK’s predictions were evaluated in test/validation sets and compared with CRISPR-spacers, VHM-net, and RaFAH. Using vHULK we obtained better general accuracy than these other programs at both the genus and species levels.

## METHODS

### Datasets

In March, 2019, we searched the Genbank nucleotide database for virus records with the following query: ‘taxid:10239 AND Sequence_type:complete’. A total of 314,536 records in format .gb were downloaded and further processed for quality control. Records without features ‘host’, ‘CDS’, ‘Translation’, or containing any format error were removed. The field “host” was checked and records without Linnaeus binomial species name as a host were also filtered out (e.g., diarrhea cat, pig, swine, chicken). Only records whose host was in domains Bacteria or Archaea were selected at this point. A total of 8,616 records passed these quality control steps, resulting in what we call *baseline dataset*.

Two predictors were generated from the baseline dataset: one for prediction of host species and another for prediction of host genus. Each of these models required its own training set; we call them the *species training set* and the *genus training set*. For adequate model training, for each host species or host genus to be predicted (i.e., to define a target), we needed minimum numbers of training instances. Thus, for a given species to be included as a target, we required the existence of at least 20 phage records with that host species; for genus, this requirement was at least 12 records. As a result, the dataset for training the genus model contained 61 targets and 7,993 instances (phage genomes), and the species model contained 52 targets for prediction and 6,432 instances (Table 1). These datasets were used for final model training (models are available with the software).

**Table 1.**
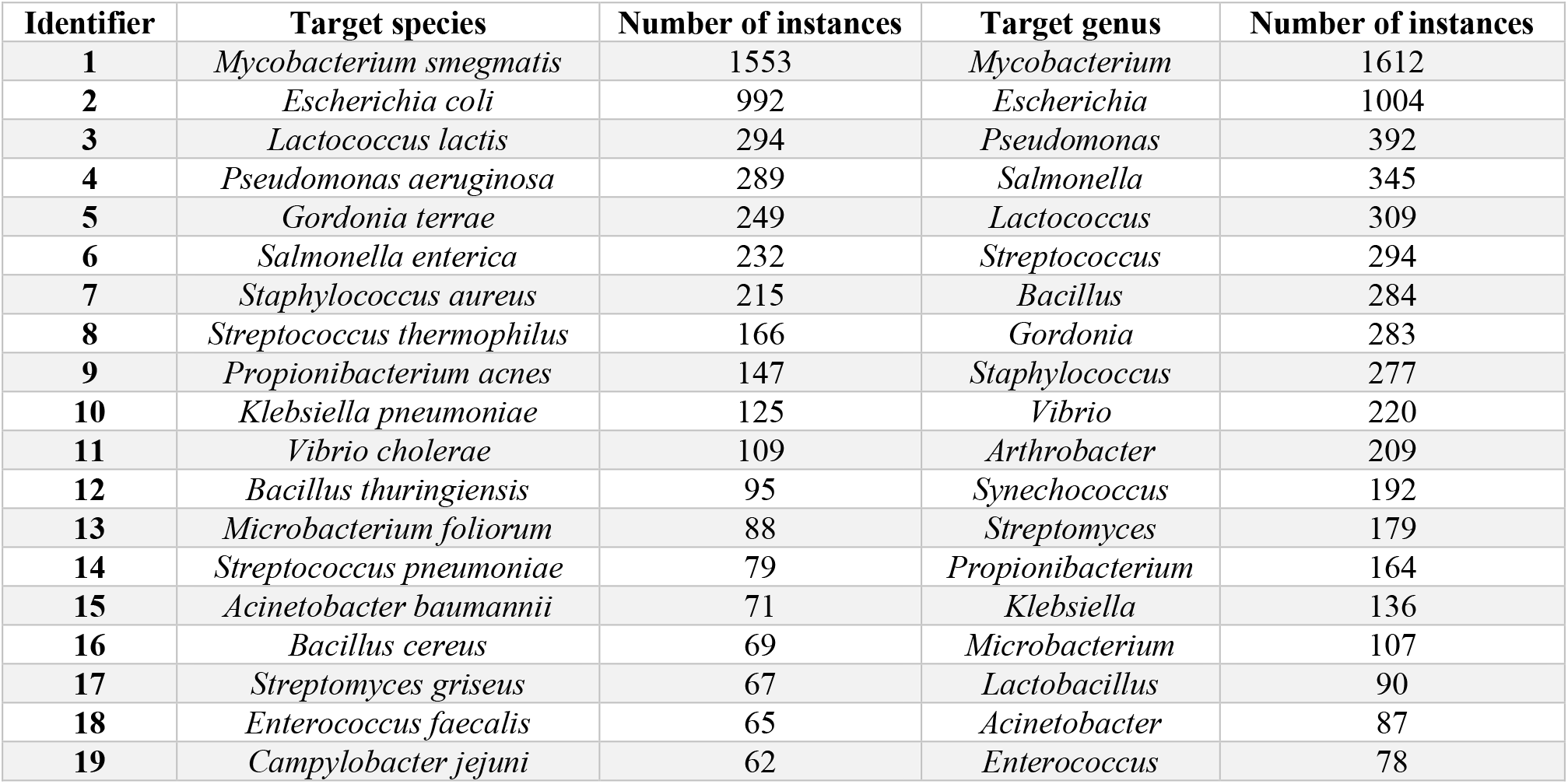
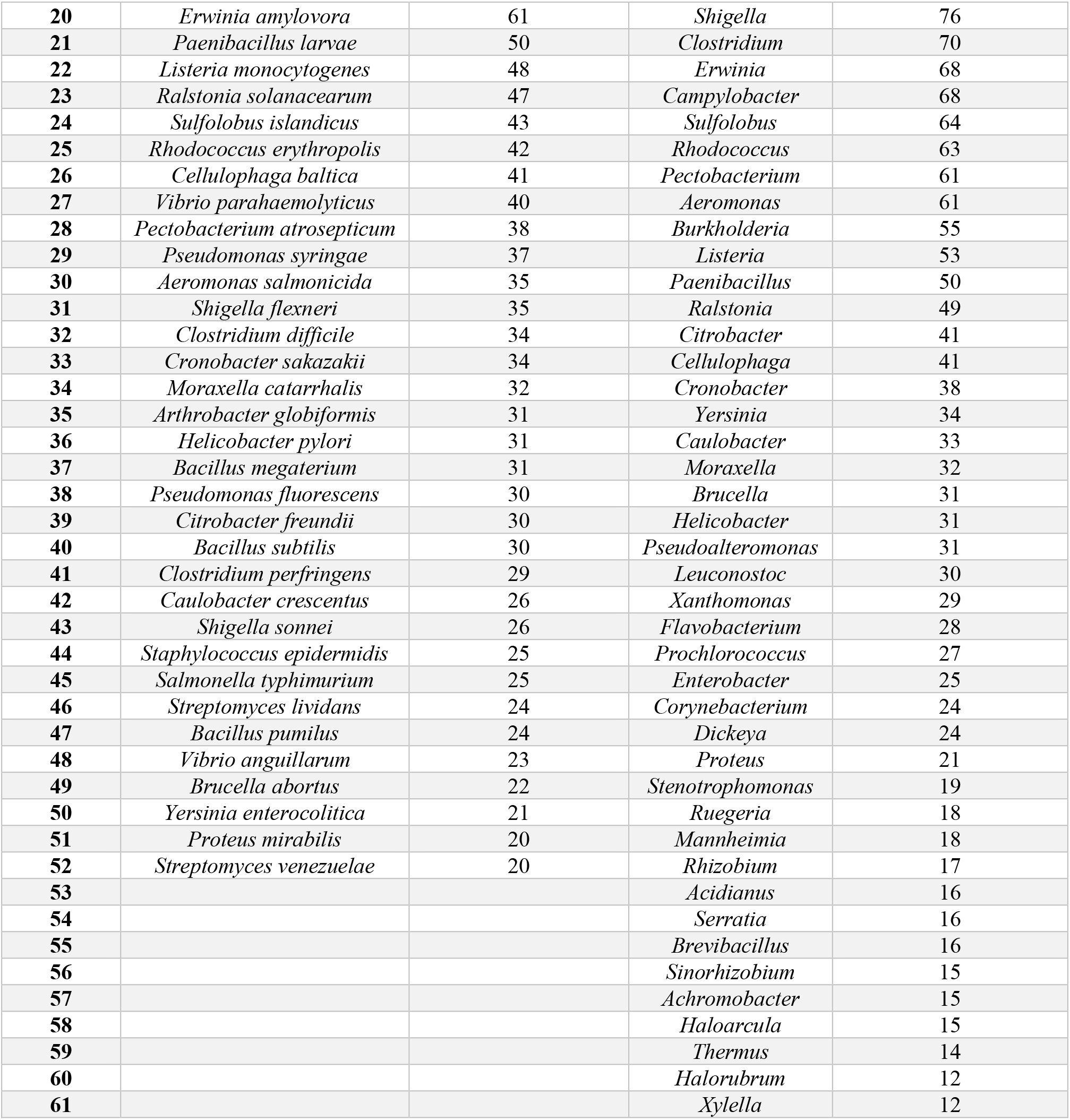
Target hosts and number of instances in the final training set for each model type (species and genus).

Because there may be very similar genomes in the species/genus training datasets, we calculated a pairwise matrix of ANI scores using the orthoANI tool (27). This matrix was used to remove redundancy from the species/genus-datasets and to generate the *performance dataset*, so that very similar genomes would not bias performance measures. Instances with ANI scores equal or higher than 97% were removed and only one randomly chosen instance from each such group was left in the performance dataset.

The performance dataset was randomly split into training and test sets in an 80/20 fashion, that is: 80% of the dataset was used to train the neural network and 20% to evaluate its predictions. Balanced accuracy was averaged for 10 random splits of the training/test datasets. Each random split test dataset had 1,414 phage genomes assessed for the genus model and 1,131 phage genomes assessed for the species model.

For validation purposes, we created the *newly-deposited-genomes (NDG) dataset*. This dataset included GenBank records released between April 1^st^, 2019 and March 31^st^, 2020, which ensured that there was no overlap with training sets. The QC filters applied to generate the baseline dataset were applied to this dataset, and complete genomes from viruses infecting any Bacteria and Archaea hosts were considered (including host targets which were not present in the training sets). A total of 2,178 genomes met these criteria.

### Feature engineering and representation

Genomes in the training species/genus datasets were individually submitted to Prokka (28) for automatic gene prediction and annotation. Predicted protein sequences (FASTA amino acid format) were searched against the pVOGs database of phage protein families (29) using HMMscan (30). Search results were parsed to populate a matrix of pVOG identifiers in columns and phage genes in rows. The matrix cell values were based on the e-value of the best alignment of each gene against the pVOGs database. We call each matrix cell value a Measure of Hit Significance (MHS), defined as follows: if e-value < 1 then MHS = 1 – e-value; if e-value ≥ 1 then MHS = 0. Values close to one mean that significant hits regarding specific phage protein families are present among phage genes for that genome. With this approach, each genome is represented by a matrix of numbers between zero and one. For efficiency reasons matrices were then transformed into vectors by adding up values in the same column; these vectors were then used as input to the neural network (Figure 1). The qualitative difference between original matrices and their vectors is that matrices also carry synteny information (relative position of a gene in the genome). We chose to use vectors instead of matrices because in preliminary testing we obtained similar performance results from both, with matrices being much more computationally intense to train.

**Figure 1.**
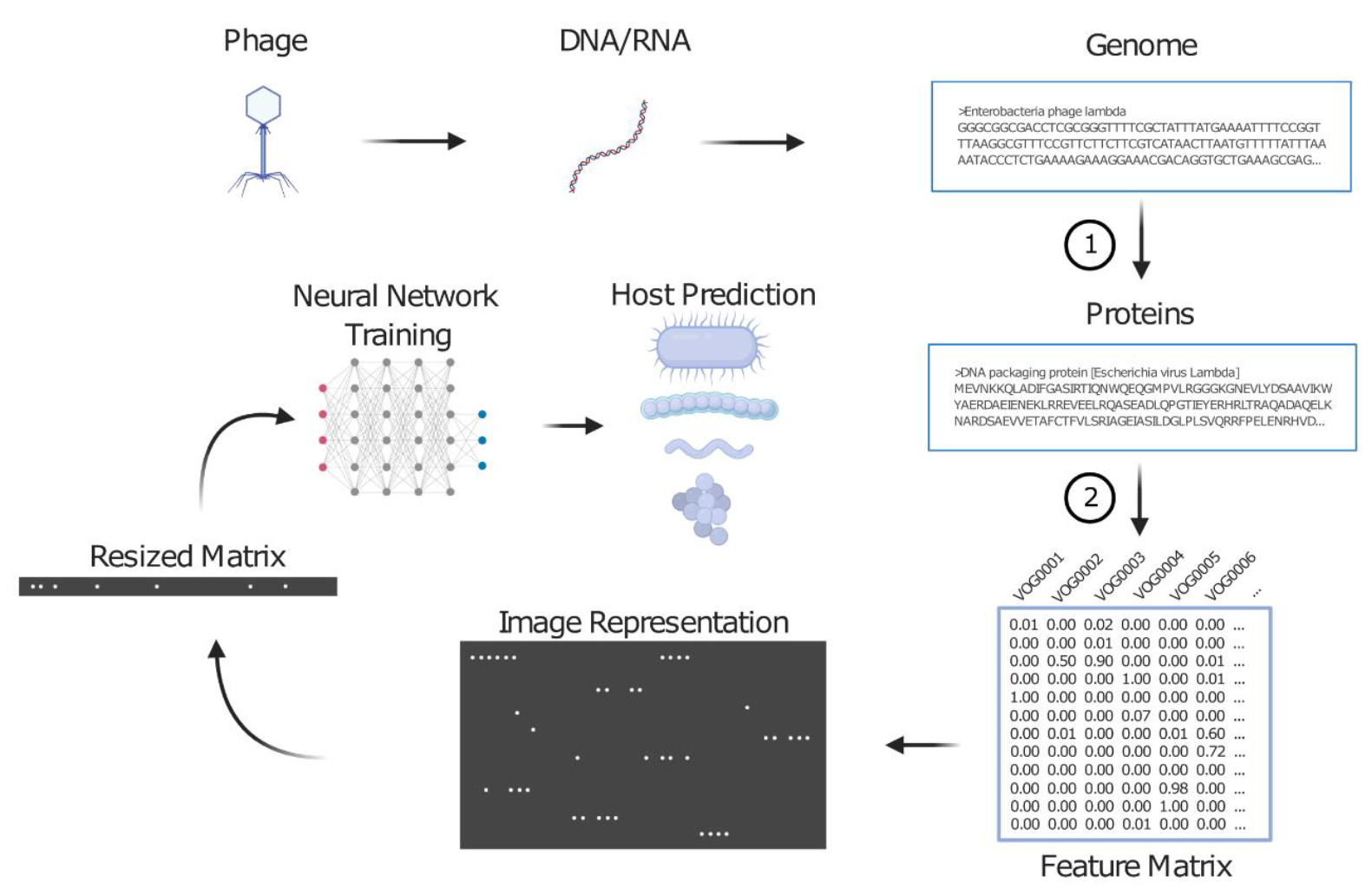
Pipeline for phage prediction. Schematic representation of how high-level features are engineered to feed the neural network and to perform host prediction. 1 - Gene prediction and annotation by Prokka; 2 - HMMscan search against pVOGs database of phage protein families and generation of a matrix with MHS values.

### Deep learning model architecture and training

Each predictor (species or genus) is composed of two internal models. Both models have the same structure except for the activation function of the last neuronal layer (31). First, we will explain the common structure and subsequently the reason for the different output functions.

Models are Multi-layer Perceptron (MLP) (32) constituted of one input layer, two hidden layers, and one output layer. The hidden layers (2nd and 3rd) have respectively 1000 and 500 neuron units with a leaky ReLU activation function. Optimization in training was achieved with the ADAM algorithm (33). Categorical cross-entropy was used as loss function with weight balancing for each host class because we had an imbalanced dataset. Early stopping was implemented during training to avoid overfitting; this allows returning the model to the last step that showed considerable decrease in the value of the loss function. The final prediction output is a vector composed of 52 prediction scores for species or 61 prediction scores for genus.

The difference between the internal models in each predictor resides on the activation function of the output layer, the layer that returns the score for each possible host. One internal model uses the ReLu activation function, which outputs values between 0 and 1. The score for a host is independent of the score for all other hosts. This means for example that all scores could be zero, which can be interpreted as meaning that the host is not among the list of possible hosts. We also implemented an internal model containing Softmax activation functions in the output layer. In this case, values for all possible predictions (which are also between 0 and 1) must add up to one, and each score may be understood as a probability. Training converges more rapidly for the Softmax internal model, and preliminary testing showed that its accuracy is slightly better. Both internal models were kept, and users can see the results of both models for each species and each genus in the list of possible species and possible genera. However, in order to provide the user with something more definite and easier to understand than just a list of scores, these outputs are put through a simple heuristic decision tree (Figure 2). It is at the end of this stage that a final, single prediction is generated. Note that a possible outcome of this decision tree is prediction ‘none’, which means that the host is not among the available targets. Even when the final prediction is ‘none’, users can still check the list of scores for both models to see whether there was some ‘good’ prediction that was left out by the heuristic thresholds of the decision tree.

**Figure 2.**
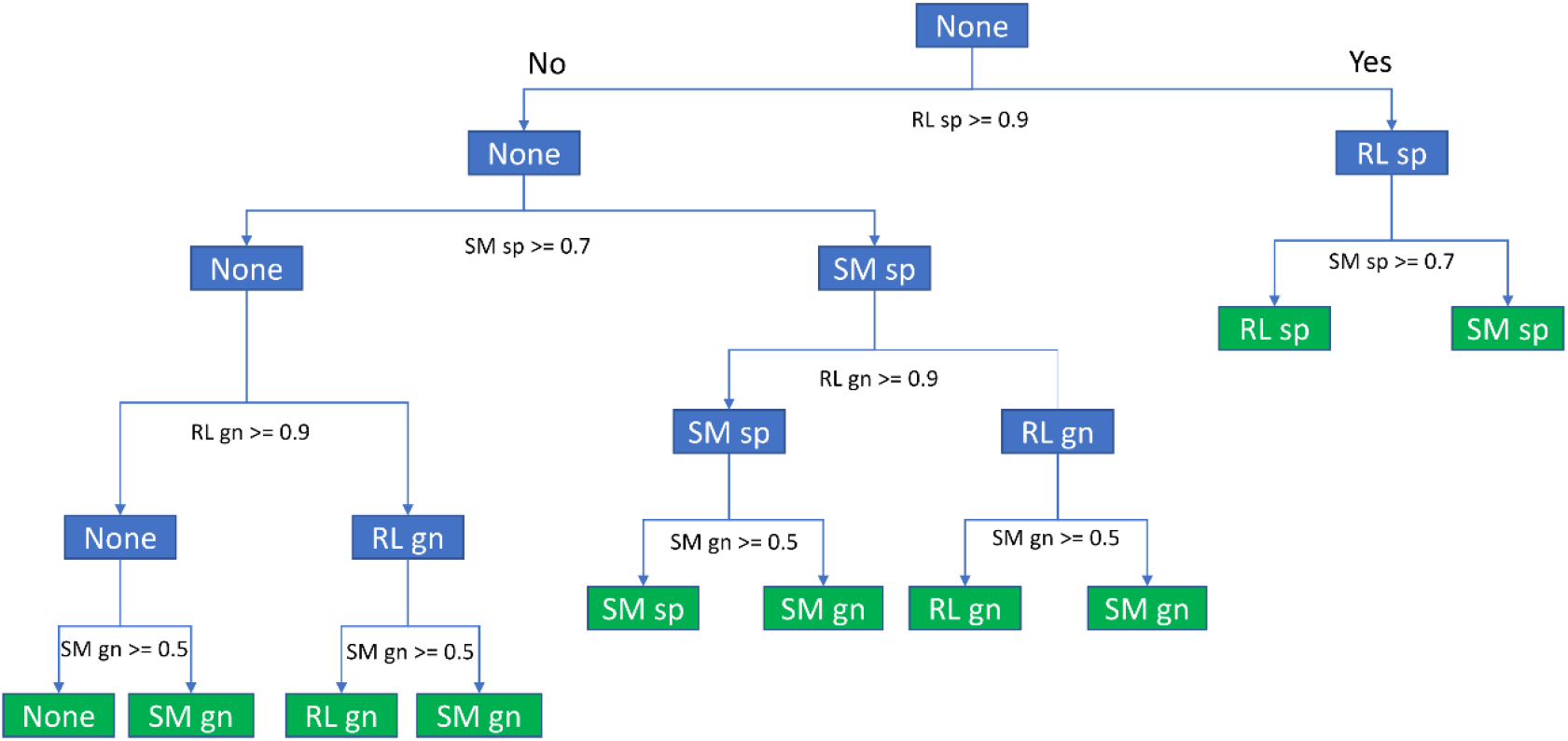
Heuristic decision tree. A decision tree was implemented to generate user-friendly and unified host predictions. RL sp represents scores for ReLu species model, RL gn for ReLu genus model, SM sp for Softmax species model and SM gn for Softmax genus model. Blue rectangular internal nodes represent intermediary decisions, and green rectangular nodes at the leaves represent the final decision reached by vHULK. The threshold values in each fork were the result of empirical observation of preliminary testing.

Final models were trained with the entire species/genus-datasets and saved as h5 files. Training code was written in Python 3 using Keras and TensorFlow libraries. Training was performed in a server equipped with a Nvidia GPU Quadros P6000 processor.

### Tool implementation

Models for prediction of host species and genus were embedded in a standalone toolkit developed in Python 3, named vHULK for Viral Host UnveiLing Kit. The tool receives phage genomes as FASTA format and outputs top prediction scores for each internal model (ReLu species, Softmax species, Relu Genus and Softmax genus). To complete the toolkit, vHULK also outputs an entropy value for the vector of prediction scores of the Softmax models, which reflects how evenly distributed are the scores for each of the target hosts. In other words, entropy will provide a measure for prediction confidence (see additional comments in the Results section). High values of entropy suggest that the model is unsure about a prediction or that a phage may infect different hosts (broad host range). Scipy implementation was used to calculate entropy from a prediction vector, as follows:

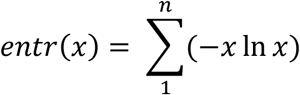

where *x* is a score in the prediction vector (0 ≤ *x* ≤ 1) and *n* is the vector size (52 for the species model and 61 for the genus model).

### Comparison of vHULK with VirHostMatcher-net, CRISPR spacers matches and RaFAH

Phage genomes of the NDG dataset were submitted for host prediction using four different programs: vHULK, neural networks with distance measurements and other features used by VirHostMatcher-net (VHM-net) (25), RaFAH (26), and CRISPR-spacers as links between host and phage genomes. For vHULK, phage genomes were individually submitted to the tool using default parameters and only the final unified prediction was considered (a score threshold was not necessary, since vHULK already makes a unified and final prediction). For VHM-net, individual genomes were submitted for prediction using default parameters. Only the best scoring hit was considered, and a threshold of at least 0.8 for the main score was applied for significant hits. Default datasets for possible hosts were used in VHM-net. For RaFAH, default parameters were used. For the CRISPR-spacers approach, individual genomes were searched against a database of CRISPR spacers of the CRISPRCasDB (34) using the NCBI blastn tool (35). To adapt our search to CRISPR spacers characteristics, word_size was set to 9 nt, and thresholds of e-value ≤ 0.001 and number of mismatches ≤ 2 were applied to select significant hits. In all four cases, predicted hosts were compared to known phage hosts as specified by the tag ‘host’ or ‘lab_host’ in the GenBank record.

## RESULTS

### Phage-host prediction features

A total of 9,504 features were generated to train the vHULK neural network models in predicting phage hosts. Features are the MHS (Measure of Hit Significance) values obtained from similarity searches to a corresponding phage orthologous group (pVOG) (29), as shown in Figure 1. Feature selection was not necessary, as neural networks perform their own feature selection by adjusting edge weights to zero (when a feature is not important for the target prediction) or to one (when it is a very informative feature). Multidimensional Scaling (MDS) representation for the entire set of features considering genus targets is shown in Figure 3. In this two-dimensional projection, some hosts appear better clustered than others. For example, the points that represent *Mycobacterium, Gordonia, Staphylococcus*, and *Streptococcus* are better clustered than those that represent *Salmonella, Escherichia*, and *Pseudomonas*. This is an indication that certain hosts may be more suitable for prediction with this set of features than others.

**Figure 3.**
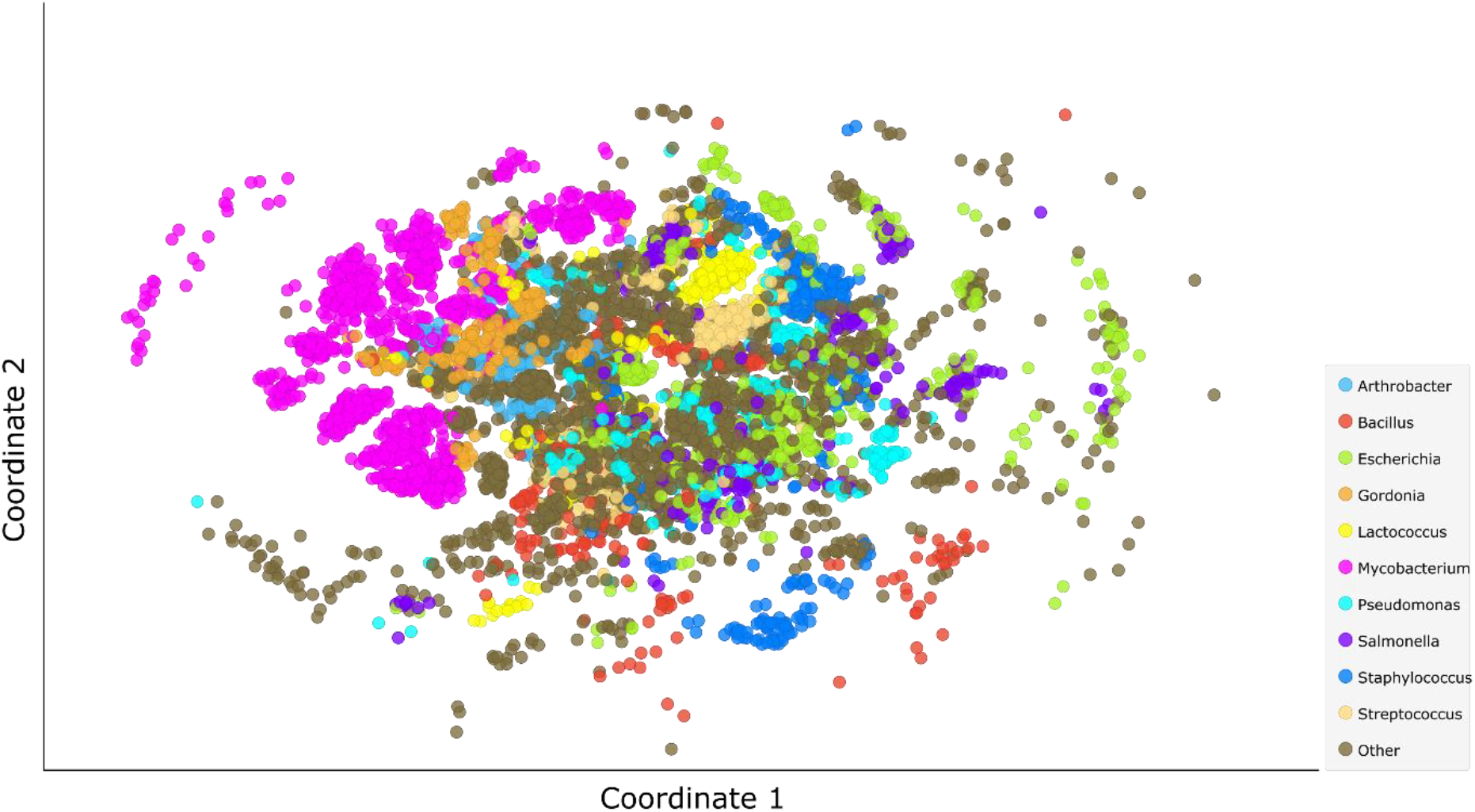
Multidimensional scaling 2D projection based on Euclidean distances of the 9,504 features used in this study. Each dot indicates a phage genome of the performance dataset. Color indicates host for the most frequent genera.

We assessed feature relevance by using a predominant correlation approach as implemented in the Fast Correlation-Based Filter algorithm (FCBF) (36). In this way we identified a set of features that seem to be strongly correlated with host prediction in specific groups of phages. Among these, features that represent the orthologous groups VOG4545, VOG4569 and VOG5417 had the highest correlation values. VOG4545 is an orthologous group ubiquitous among the three tailed phages families (*Myoviridae*, *Siphoviridae* and *Podoviridae*), composed of 1,747 known orthologous proteins. Most of the VOG4545 member proteins are annotated as a tape measure protein, which is thought to vary in length according to phage tail size (37). Proteins of the orthologous group VOG4569 are present in phages from families *Siphoviridae* and *Podoviridae*, mainly infecting Actinobacteria (*Mycobacterium* and *Gordonia*). They are annotated as a structural minor tail protein. Lastly, the orthologous group VOG5417 is composed of proteins annotated as a lysin B. Members of the group are present in phages that infect Actinobacteria. It is worth noting that the pVOG annotations of all three best correlated families indicate that these proteins are directly or indirectly related to the infection mechanism. Correlation and information theory metrics for all features used in this study are available as Supplementary Table S1.

### Neural Network Training and Testing

We monitored learning curves and the cross-entropy loss function during model training. Results indicate that models quickly converged to a plateau of accuracy, with Softmax internal models performing slightly better than ReLu internal models (Figure 4). Maximum performance was reached around the second epoch iteration in all training sets. No more than six epochs were necessary for triggering the early stop algorithm and finish training. Training and validation accuracies did not greatly diverge in values, suggesting that overfitting was not a problem during training.

**Figure 4.**
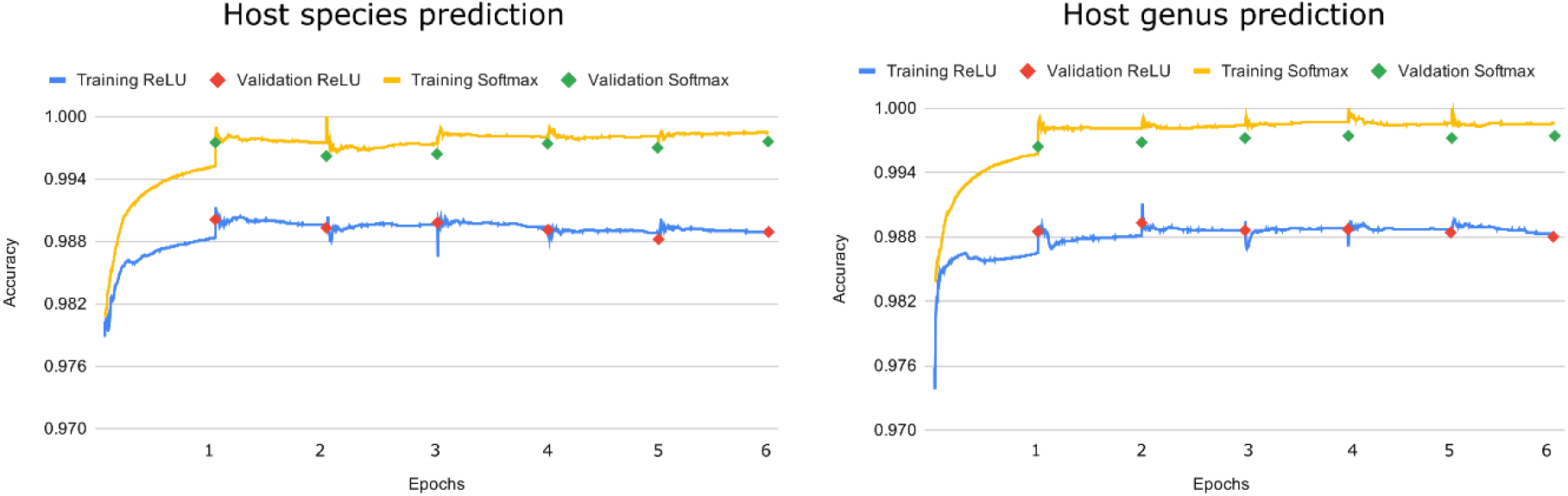
Training accuracy using the performance dataset for the four internal models developed in this study. X-axis shows epoch iteration, i.e., points where the training algorithm passed through the entire dataset. Validation in this figure means using the testing dataset to evaluate performance after each epoch is completed.

On the nonredundant test datasets (put aside from training sets), average accuracy for ten random splits was 99.1% for predictions at the genus level and 98.9% for predictions at the species level (Table 2).

**Table 2.** Average accuracy and standard deviation for ten random splits of the performance dataset

|  | Average Accuracy | Standard deviation |
| --- | --- | --- |
| Prediction at species level<sup>a</sup> | 0.98983 | 0.00172 |
| Prediction at genus level<sup>a</sup> | 0.99129 | 0.00076 |

<sup>a</sup> Genomes in the testing set were not used to train the model.

A confusion matrix for predictions at the genus level for one of the random split test sets is shown by heatmaps in Figure 5. Hierarchical clustering (Figure 5B) shows that most of the confusion happened with hosts from a same taxonomic family (clusters highlighted in gray). These results are expected and may be biologically meaningful, since wide host-range phages have been reported to infect different genera from the same family (38,39).

**Figure 5.**
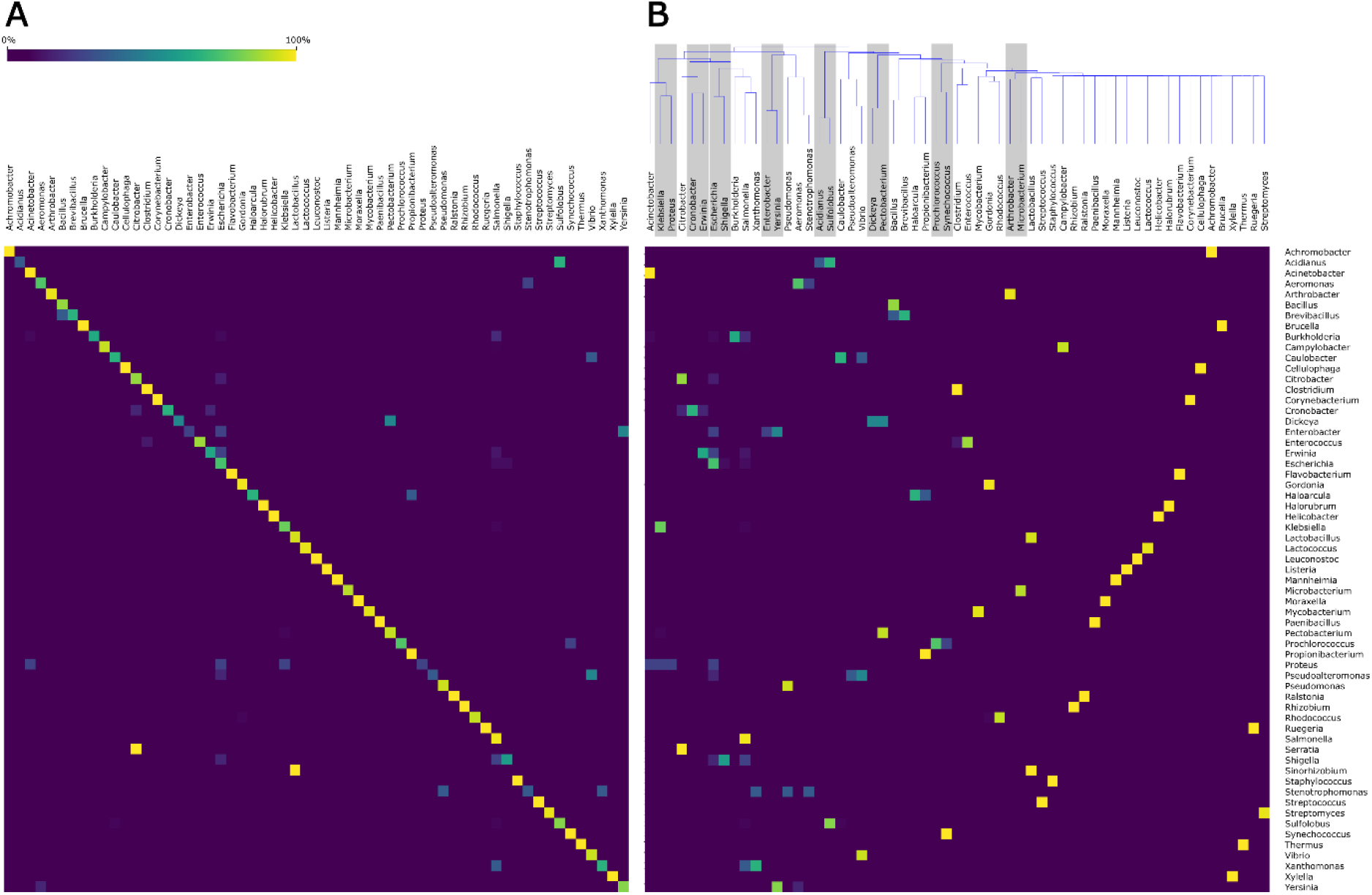
Confusion matrices of host predictions. Predictions shown are for genus for one of the random split test datasets. Confusion matrices are shown as heatmaps, where values correspond to the percentage of each actual host (matrix rows) that was predicted as a specific host (matrix columns). (**A**) Hosts are listed in alphabetical order; points on the diagonal are correct predictions. (**B**) The same predictions as in (**A**) are shown hierarchically clustered by frequent confusions. Hosts highlighted by a grey box belong to the same taxonomical family.

### Validation with the Newly Deposited Genomes dataset

We used the *newly-deposited-genomes* (*NDG*) *dataset* for validation purposes (see Methods). Performance on the NDG dataset varied depending on the host and type of prediction (genus or species level). Table 3 shows performance results for the 10 most frequent host species present in the dataset. Accuracy was better than 96% at both genus and species levels for 9 out of 10 hosts; it was 93% for phages infecting *Escherichia coli*. Specificity was greater than 97% for all hosts at both taxonomic levels. For six hosts, recall rates were 80% or better at both levels. Particularly low recall values were observed for *Salmonella enterica* (genus: 50%, and species: 35.4%) and *Klebsiella pneumoniae* (genus: 60.95%, and species: 55.8%). Average entropy values for each of the top 10 hosts are also shown in Table 3. Predictions for Salmonella- and Klebsiella-infecting phages were the only ones in which entropy was higher than 1.

**Table 3.** Summary of vHULK performance results for the 10 most frequent species in the NDG dataset.

| Genus/Species | Count genus | Count species | Genus prediction |  |  | Species prediction |  |  | Most frequent wrong prediction<sup>a</sup> | Entropy species | Entropy genus |
| --- | --- | --- | --- | --- | --- | --- | --- | --- | --- | --- | --- |
|  |  |  | Recall | Specificity | Accuracy | Recall | Specificity | Accuracy |  |  |  |
| *Mycobacterium smegmatis* | 332 | 330 | 100.00 | 99.13 | 99.26 | 100.00 | 99.45 | 99.54 | - | 0.0009 | 0.009 |
| *Escherichia coli* | 265 | 265 | 81.88 | 97.07 | 95.22 | 54.33 | 98.43 | 93.06 | *Campylobacter jejuni* (29) | 0.5548 | 0.5548 |
| *Gordonia terrae* | 151 | 122 | 90.06 | 99.80 | 99.12 | 94.26 | 98.88 | 98.62 | *Mycobacterium* (14) | 0.1958 | 0.2801 |
| *Salmonella enterica* | 114 | 113 | 50.00 | 99.51 | 97.38 | 35.39 | 99.61 | 96.28 | *Escherichia* (27) | 1.06 | 1.0598 |
| *Pseudomonas aeruginosa* | 116 | 103 | 88.79 | 99.51 | 98.94 | 92.23 | 99.75 | 99.40 | None (7) | 0.24 | 0.3359 |
| *Microbacterium foliorum* | 115 | 98 | 87.82 | 100.00 | 99.35 | 87.75 | 99.37 | 98.85 | None (9) | 0.464 | 0.4921 |
| *Klebsiella pneumoniae* | 105 | 86 | 60.95 | 99.80 | 97.93 | 55.81 | 99.23 | 97.52 | *Erwinia amylovora* (10) | 1.1553 | 1.1212 |
| *Streptococcus pyogenes*<sup>b</sup> | 330 | 67 | 84.24 | 100.00 | 97.61 | - | - | - | None (51) | 0.171 | 0.6117 |
| *Lactococcus lactis* | 55 | 55 | 100.00 | 100.00 | 100.00 | 100.00 | 100.00 | 100.00 | - | 0.0596 | 0.0596 |
| *Enterococcus faecalis* | 40 | 40 | 97.50 | 100.00 | 99.95 | 70.00 | 100.00 | 99.44 | None (1) | 0.097 | 0.097 |
| *Acinetobacter baumannii* | 41 | 38 | 97.56 | 99.9 | 99.86 | 94.73 | 99.85 | 99.77 | None (1) | 0.08 | 0.1368 |

<sup>a</sup> Most frequent wrong prediction is indicated with an absolute count between parentheses.
<sup>b</sup> Predictions at the species level were not assessed for *Streptococcus pyogenes* because it is not one of the target hosts in vHULK species models.

It is worth noting that entropy is generally higher for hosts whose prediction accuracy is lower. To assess the hypothesis of a significant linear correlation between accuracy and entropy, we calculated Spearman’s correlation coefficient for the ten most frequent hosts in the NDG dataset. Results show a coefficient Rs = −0.879 (p-value < 0.001) for predictions at the species level and Rs = −0.782 (p-value < 0.001) for predictions at the genus level. This is a relevant information to vHULK users, who may use entropy outputs as additional information and a proxy for confidence in predictions. Each model’s individual predictions, along with scores and entropy for all phage genomes in the NDG dataset, are available as Supplementary Table S2.

### Comparing host prediction programs

We have compared vHULK host predictions to those provided by VHM-net, RaFAH, and CRISPR-spacers matches. The NDG dataset was used for this purpose. As shown in Figure 6, vHULK mean accuracy (across all hosts) was 4 times higher than the second-best approach when predictions at the species level are considered (52% against 13% of CRISPR-spacers). When predictions at the genus level are considered, vHULK obtained a mean accuracy of 81.8% across 61 possible host targets, while RaFAH (the second best) obtained 71.3%. Detailed information about predictions in the NDG dataset for the four methods is available in Supplementary Table S3.

**Figure 6.**
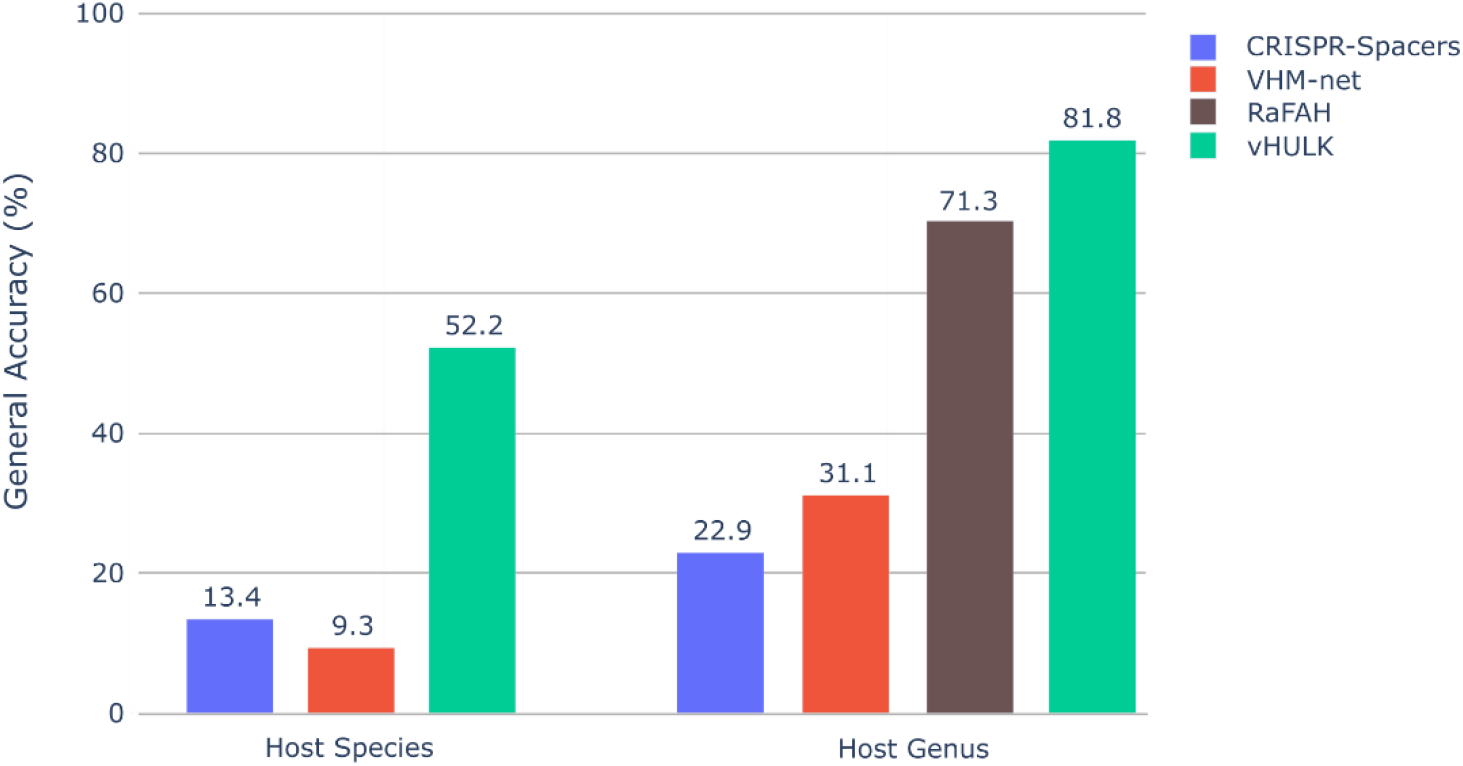
General average accuracy using vHULK, VirHostMatcher-net RaFAH tools, as well as the CRISPR-spacers matches approach. Predictions were performed for 2,178 phage genomes of the NDG dataset, both at the species and genus levels.

All four programs ran on a server with Intel Xeon 2 GHz CPUs, 22 threads and 32 GB of RAM. vHULK took approximately 18 h of wall time, VHM-net took approximately 32 h, CRISPR-spacers took approximately 50 minutes, and RaFAH took approximately 15 h. Genomes were submitted sequentially, and no additional parallelization was used in any of the tools.

## DISCUSSION

We have presented vHULK, a new program to predict phage hosts based on genome sequences. Testing showed that vHULK achieved 99% of average accuracy for both host genus and species. Performance on a validation set with 2,178 genomes that were not part of training and testing (the NDG dataset) resulted in lower accuracy values than those observed in testing (52% for species and 82% for genus). On the other hand, the three other methods that we tested had worse results on the NDG dataset than did vHULK, suggesting that this dataset is currently a difficult test for viral host prediction methods.

The noted difference in vHULK’s performance (primarily at the species level) between testing and validation could be explained in part by the presence of phages in the NDG dataset whose hosts are not present in vHULK’s training datasets. It is known that many machine learning prediction models may struggle when facing instances for which there was no target in training (24,25,40). We have tried to mitigate this problem by providing users with entropy values associated with scores; we have shown that these entropy values can be used as proxies for prediction confidence.

vHULK was trained to predict 52 different species and 61 different genera. These numbers of possible hosts were limited only by the number of instances available to train the model and by our requirement that there had to be at least 20 records for any given species and at least 12 records for any given genus. One can ask whether these 52 species and 61 genera are representative of currently available data. We can answer this question in part by noting that the 52 species account for 70% of all host species present in the NDG dataset, and the 61 genera account for 97% of all host genera present in the NDG dataset. On the other hand, 52 and 61 are tiny numbers when compared with the presumed richness of phage hosts that exist in the entire biosphere. We expect however that the number of species and genus hosts in the vHULK repertoire will grow with future releases of the software.

Despite this relatively small host repertoire, we envision that vHULK will have wide application. One use could be the mining of metagenomic datasets for new phages that may have as host one of the six multidrug resistant bacterial pathogens that compose the ESKAPE group (*Enterococcus faecium, Staphylococcus aureus, Klebsiella pneumoniae, Acinetobacter baumannii, Pseudomonas aeruginosa, Enterobacter spp*.) (41). All of them are part of the vHULK host repertoire.

Regarding the design of vHULK, we would like to point out that the program embodies a novel approach for extracting and representing genomic features for use in deep neural networks. The main novelty is the use of alignment significance scores (MHS) when evaluating presence/absence of protein families in phage genomes as features in the neural network. Our results demonstrate that the use of these features was a determining factor in achieving good accuracy. We conjecture that this same approach can be extended to the prediction of other viral biological attributes, such as viral life cycle and virulence.

F. Young and colleagues have recently evaluated different levels of genomic features for phage host prediction (42). These authors defined four levels of features that could be extracted from genomes by applying different annotation processes: Oligonucleotide frequencies, amino acid frequencies, protein physico-chemical properties, and protein domains. They conclude that all feature levels yield significant signals for host prediction, probably being complementary to one another. The protein features in vHULK are somewhat related to the protein domains of Young et al. (42). However, whereas they proposed a count of domains, we use alignment significance scores. To our knowledge, this type of machine learning feature has never been used in this application.

In sum, our results show that vHULK achieves the best performance among existing programs for phage host prediction. We hope that its availability will be found useful by the research community interested in such predictions.

## SOFTWARE AVAILABILITY

The tool, as well as testing examples and documentation is publicly available at GitHub: https://github.com/LaboratorioBioinformatica/vHULK. We also provide GenBank records used to train vHULK’s models and accessions numbers for the NDG dataset in http://projetos.lbi.iq.usp.br/phaghost/vHULK/datasets.

## ACKNOWLEDGMENTS

The Nvidia GPU Quadro P6000 unit used in this work was obtained through a donation by the Nvidia Corporation to JCS.

